# SARS-CoV-2 receptor ACE2 identifies immuno-hot tumors in breast cancer

**DOI:** 10.1101/2021.05.10.443377

**Authors:** Jie Mei, Yun Cai, Rui Xu, Xinqian Yu, Lingyan Chen, Tao Ma, Tianshu Gao, Fei Gao, Yichao Zhu, Yan Zhang

## Abstract

Angiotensin-converting enzyme 2 (ACE2) is known as a host cell receptor for Severe Acute Respiratory Syndrome Coronavirus 2 (SARS-CoV-2), which is identified to be dysregulated in multiple tumors. Although the characterization of abnormal ACE2 expression in malignancies has been preliminarily explored, in-depth analysis of ACE2 in breast cancer (BRCA) has not been elucidated. A systematic pan-cancer analysis was conducted to assess the expression pattern and immunological role of ACE2 based on RNA-sequencing (RNA-seq) data downloaded from The Cancer Genome Atlas (TCGA). Next, correlations between ACE2 expression immunological characteristics in the BRCA tumor microenvironment (TME) were evaluated. Also, the role of ACE2 in predicting the clinical features and the response to therapeutic options in BRCA was estimated. These findings were subsequently validated in another public transcriptomic cohort as well as a recruited cohort. ACE2 was lowly expressed in most cancers compared with adjacent tissues. ACE2 was positively correlated with immunomodulators, tumor-infiltrating immune cells (TIICs), cancer immunity cycles, immune checkpoints, and tumor mutation burden (TMB). Besides, high ACE2 levels indicated the triple-negative breast cancer (TNBC) subtype of BRCA, lower response to endocrine therapy and higher response to chemotherapy, anti-ERBB therapy, antiangiogenic therapy and immunotherapy. To sum up, ACE2 correlates with an inflamed TME and identifies immuno-hot tumors, which may be used as an auxiliary biomarker for the identification of immunological characteristics in BRCA.

## Introduction

Coronavirus disease 2019 (COVID-19) is infectious pneumonia caused by Severe Acute Respiratory Syndrome Coronavirus 2 (SARS-CoV-2) infection [1]. At the end of 2019, COVID-19 was originally reported in China in December 2019. Just over a year, COVID-19 has transmitted fastly to almost all countries, which leads to a serious global public health problem. Based on the latest statistics released by World Health Organization, a total of 146,054,107 confirmed cases and 3,092,410 deaths have been reported as of 25 April, 2021 (https://covid19.who.int/). Angiotensin-converting enzyme 2 (ACE2) is recognized as a host cell receptor for SARS-CoV and SARS-CoV-2 [2, 3]. Emerging research reported that the expression and distribution of ACE2 were tissue-specific to some extent, which is enriched in the lung, esophagus, kidney, bladder, testis, stomach and ileum using a single-cell RNA sequencing (RNA-seq) technique [4]. However, ACE2 is expressed in all organs, excepting for the prostate and brain, although some organs exhibit low expression [5]. Thus, organs expressing high ACE2 appear to be more impressionable to SARS-CoV-2 infection in healthy individuals.

Breast cancer (BRCA) is a multifactorial disease, which has the highest incidence in the world. In 2020, a total of 2,261,419 new cases and 684,996 deaths have been reported according to the latest statistics [6]. Be a multifactorial disease, dysregulation of the immune landscape acts as a significant role in the oncogenesis and development of BRCA, which lays the molecular foundation for immunotherapy [7]. ACE2 is known as a tumor suppressor and is lowly expressed in most cancers [8-10]. Encouragingly, several analyses reveal that ACE2 correlates with the abundances of a number of tumor-infiltrating immune cells (TIICs) in multiple cancers [11, 12]. However, indiscriminate pan-cancer analysis neglects in-depth research on dominant tumor species, which may lead to ignoring the great value of ACE2 in regulating tumor immunity and acting as a indicator for the stratification of tumor immunogenicity.

Tumors are complex masses consisting of malignant as well as normal cells. The multiple interplays between these cells via cytokines, chemokines and growth factors constitute the tumor microenvironment (TME) [13]. TME could be crucial for the response to several therapies and the prognosis. Tumors can be simply classified into cold or hot depending on their TME. Cold tumors are characterized as immunosuppressive TME and insensitive to either chemotherapy or immunotherapy, and hot tumors represent higher response rates to these therapies, which is featured by T cell infiltration and immunosuppressive TME [14]. In principle, the hot tumors exhibit a good response to immunotherapy, such as anti-PD-1/PD-L1 therapy [15]. Thus, distinguishing hot and cold tumors is a critical strategy to demarcate the response to immunotherapy.

In this research, we first conducted a pan-cancer analysis of the expression and immunological features of ACE2. We discovered that ACE2 exhibited the tightest correlation with immunological factors in BRCA, which may be a dominant tumor species for the in-depth analysis of the immunological role of ACE2. We also revealed that ACE2 indicated an inflamed TME and identified immuno-hot tumors in BRCA, and had the potential to estimate the molecular subtype of BRCA.

## Methodology

### Public datasets retrieval

The Cancer Genome Atlas (TCGA) data: The pan-cancer normalized RNA-seq datasets, copy number variant (CNV) data processed by GISTIC algorithm, 450K methylation data, mutation profiles, the activities of transcription factor (TF) calculated by RABIT, and clinical information were obtained from UCSC Xena data portal (https://xenabrowser.net/datapages/). The somatic mutation data were obtained from TCGA (http://cancergenome.nih.gov/) and then used to calculate the tumor mutation burden (TMB) by R package “maftools”. The abbreviations for TCGA cancer types are shown in Table S1.

METABRIC data: The normalized RNA-seq dataset, CNV data processed by GISTIC algorithm, mutation profiles and clinical data of BRCA patients in METABRIC cohort were downloaded from cBioPortal data portal (http://www.cbioportal.org/datasets) [16].

### Prognostic analysis using PrognoScan

PrognoScan database (http://dna00.bio.kyutech.ac.jp/PrognoScan/) was applied to assess the prognostic value of ACE2 in BRCA across a large cohort of public microarray datasets [17]. All the results were exhibited in the study.

### Linked Omics database analysis

The Linked Omics database (http://www.linkedomics.org/login.php) is a web-based tool to analyze multi-dimensional datasets [18]. The functional roles of ACE2 in BRCA was predicted using the Linked Omics tool in term of Gene Ontology (GO) and Kyoto Encyclopedia of Genes and Genomes (KEGG) pathways by the gene set enrichment analysis (GSEA). Default options were used for all parameters.

### Assessment of the immunological features in TME of BRCA

The immunological features of TME in BRCA contain immunomodulators, the activities of the cancer immunity cycle, infiltration levels of TIICs, and the expression of inhibitory immune checkpoints.

The information of 122 immunomodulators including major histocompatibility complex (MHC), receptors, chemokines, and immunostimulators was collected from the research of Charoentong et al. [19]. Considering the cancer immunity cycle which contains seven stages reflects the anti-cancer immune response and the activities of each step decide the fate of tumor cells, we subsequently calculated the activation scores of each step by single sample gene set enrichment analysis (ssGSEA) according to the expression level of specific signatures of each step [20]. Moreover, in order to avoid calculation errors resulted from various algorithms which were developed to explore the relative abundance of TIICs in TME, we comprehensively estimated the infiltration levels of TIICs using the following independent algorithms: TIMER [21], EPIC [22], MCP-counter [23], quanTIseq [24] and TISIDB [25]. ESTIMATE algorithm was also performed to calculate Tumor Purity, ESTIMATE Score, Immune Score and Stromal Score [26]. Furthermore, we also collected several well-known effector genes of TIICs, and computed the T cell inflamed score according to the expression levels and weighting coefficient of 18 genes reported by Ayer et al. [27].

To verify the role of ACE2 in mediating cancer immunity in BRCA, we grouped the patients into the high ACE2 and the low ACE2 group with the 50% cutoff based on the expression levels of ACE2, and then analyzed the difference of the immunological features of TME concerning the above aspects between the high and low ACE2 groups.

### Immunophenoscore analysis

As previously reported, a patient’s immunophenoscore (IPS) can be calculated without bias using machine learning by consideration of the 4 major categories of components that measure immunogenicity: effector cells, immunosuppressive cells, MHC molecules, and immunomodulators [19]. The IPS values of BRCA patients were obtained from the Cancer Immunome Atlas (TCIA) (https://tcia.at/home).

### Calculation of the enrichment scores of various gene signatures

According to previous research [28], we collected several gene-sets positively associated with therapeutic response to immunotherapy, targeted therapy and radiotherapy, and specific markers of biological process correlated with anti-tumor immunity such as genes involved in DNA replication. The enrichment scores of these signatures were obtained using the GSVA R package [29].

### Prediction of therapeutic response

The role of ACE in predicting the response to chemotherapy was also evaluated. First, BRCA-related drug-target genes were screened using the Drugbank database (https://go.drugbank.com/). Besides, we predicted the response to anti-therapy for each patient based on the Cancer Genome Project (CGP) database (https://www.sciencedirect.com/topics/neuroscience/cancer-genome-project) as well. Several common therapeutics, including vinblastine, cisplatin, doxorubicin, etoposide, gefitinib, gemcitabine, paclitaxel, parthenolide, sunitinib and vinorelbine were selected. The prediction process was performed by R package “pRRophetic” where the samples’ half-maximal inhibitory concentration (IC50) was calculated by ridge regression and the prediction accuracy was assessed by 10-fold cross-validation according to the CGP training set. Default options were used for all parameters [30].

### Clinical samples

Two tissue microarrays (TMAs, HBreD050Bc01 and HBreD090Bc03) were obtained from Outdo Biotech (Shanghai, China). The HBreD050Bc01 microarray contained 40 BRCA and 10 adjacent samples. The HBreD090Bc03 microarray contained 85 BRCA and 5 adjacent samples. Thus, a total of 125 BRCA samples and 15 adjacent samples were involved in the current research.

### Immunohistochemistry and semi-quantitative evaluation

Next, Immunohistochemistry (IHC) staining was conducted on these tissue slides. The primary antibodies used in the research were as following: anti-ACE2 (1:3000 dilution, Cat. ab15348, Abcam, Cambridge, UK), anti-CD8 (Ready-to-use, Cat. PA067, Abcarta, Suzhou, China) and anti-PD-L1 (Ready-to-use, Cat. GT2280, GeneTech, Shanghai, China). Antibody staining was visualized with DAB and hematoxylin counterstain, and stained sections were scanned using Aperio Digital Pathology Slide Scanners. All stained sections were independently evaluated by two independent pathologists. For semi-quantitative evaluation of ACE2 and PD-L1 staining, the percentage of positively stained cells was scored as 0-4: 0 (< 1%), 1 (1-5%), 2 (6-25%), 3 (26-50%) and 4 (> 50%). The staining intensity was scored as 0-3: 0 (negative), 1 (weak), 2 (moderate) and 3 (strong). The immunoreactivity score (IRS) equals to the percentages of positive cells multiplied with staining intensity. For CD8 staining, infiltration level was assessed by estimating the percentage of cells with strong intensity of membrane staining in the stroma cells.

### Statistical analysis

Statistical analysis and figure exhibition performed using R language 4.0.0. The statistical difference of continuous variables between the two groups was evaluated by Wilcoxon rank sum test or Mann-Whitney test and chi-square test was used when the categorical variables were assessed. Pearson’s correlation was used to evaluete the correlation between two variables. Receiver-operating characteristic (ROC) analysis was plotted to assess the specificity and sensitivity of the candidate indicator, and the area under the ROC curve (AUC) was generated for diagnostic biomarkers. Prognostic values of categorical variables were assessed by log-rank test. For all analyses, P value ≤ 0.05 were deemed to be statistically significant.

## Results

### Expression and immunological roles of ACE2 across human cancers

After a systematic pan-cancer analysis of the expression of ACE2 in the TCGA database, we discovered that ACE2 was lowly expressed in a fraction of cancers, including BRCA, KICH, and LUAD. However, ACE2 also shown to be overexpressed in CESE, KIRP, LIHC and UCEC (Figure 1A). Next, we conducted a pan-cancer survival analysis about overall survival (OS), progression-free survival (PFS) using Kaplan-Meier analysis. ACE2 emerged as a prognostic risk factor for both OS and PFS in KIRC, LIHC and UCS (Figure S1A-B). However, these results need further verification, especially based on recruited cohorts.

**Figure 1.**
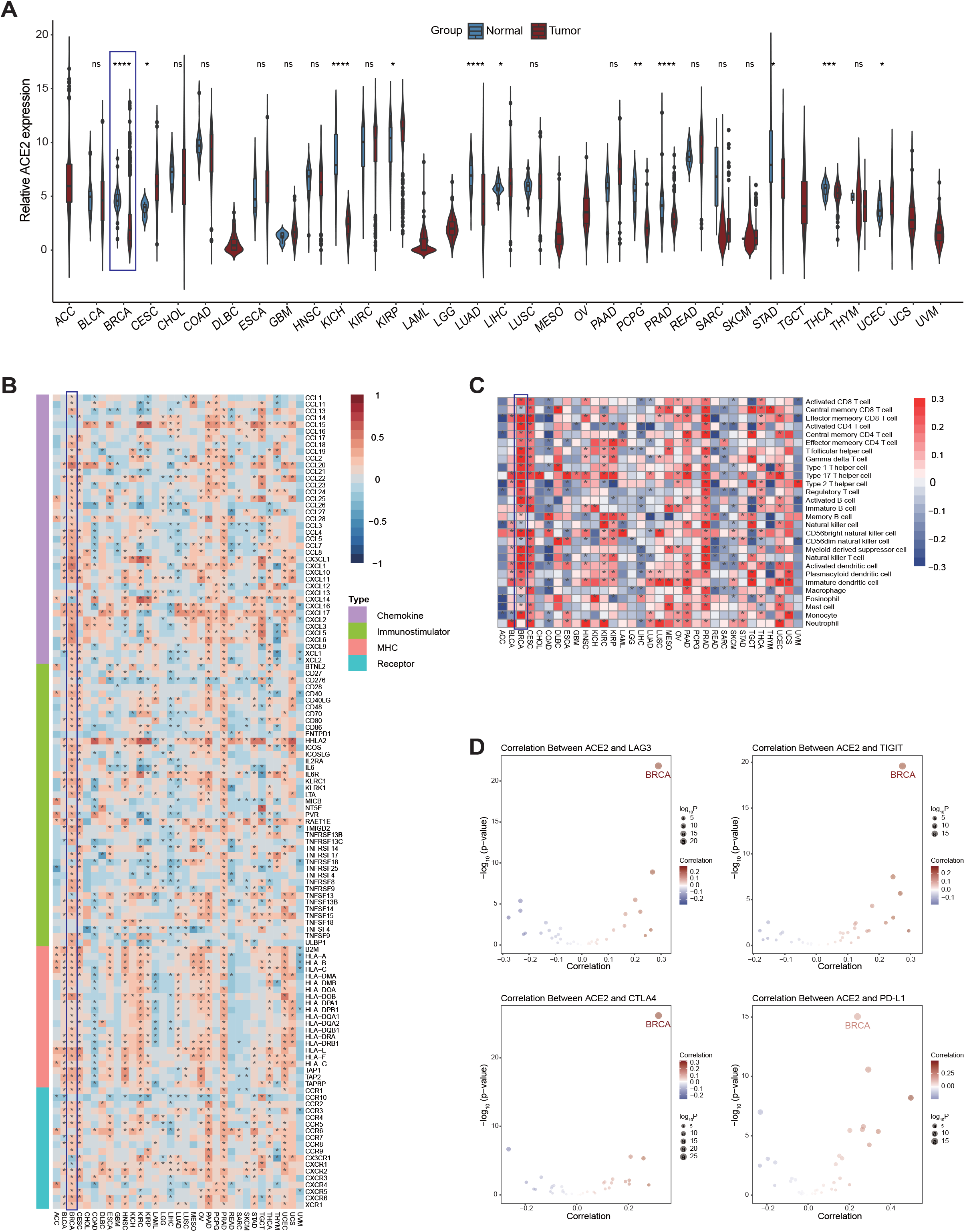
The relationship between ACE2 and immunological features in pan-cancers (A) Pan-cancer analysis of ACE2 expression in tumor and paracancerous tissues in the TCGA database. Ns: no significant difference; *P < 0.05; **P < 0.01; ***P < 0.001; ****P < 0.0001. (B) Correlation between ACE2 and 122 immunomodulators (chemokines, receptors, MHC and immunostimulators). The color reveals the correlation coefficient. The asterisks reveal statistical difference assessed by Pearson analysis. (C) Correlation between ACE2 and 28 TIICs calculated with the ssGSEA algorithm. The color reveals the correlation coefficient. The asterisks reveal statistical difference assessed by Pearson analysis. (D) Correlation between ACE2 and four immune checkpoints, LAG3, TIGIT, CTLA4, PD-L1. The dots symbolize cancer types.

We next conducted a pan-cancer analysis aimed to examine the immunological features of ACE2 in various tumors. The results uncovered that ACE2 was positively correlated with most immunomodulators in BRCA (Figure 1B). We also calculated the infiltrating levels of TIICs in the TME by the ssGSEA method. Similarly, ACE2 was positively correlated with most TIICs in BRCA (Figure 1C). Moreover, we revealed that the expression of ACE2 was positively related to the expression of several immune checkpoints, including LAG3, TIGIT, CTLA4 and PD-L1 in BRCA (Figure 1D). Although these positive correlations between ACE2 and immunological features were found in other tumors, such as CESC, KIRC and PRAD, the highest correlation was observed in BRCA. Moreover, according to previous research, ACE2 was not expressed in immune cells, and the expression of ACE2 in bulk RNA-seq data was derived from non-immune cells in all probability, such as tumor cells in the tissues.[31] Besides, considering the positive correlation between ACE2 and PD-L1, an immune checkpoint expressed on tumor cells, we conducted the TFs network analysis, and found that a mass of shared TFs that potentially regulated ACE2 and PD-L1 (Figure S2, Table S2). Collectively, the expression pattern of ACE2 is TME- characteristic, which illustrates the potential of ACE2 as an immune-related biomarker and therapeutic target in BRCA.

### Potential regulatory factors, prognostic value and functions of ACE2 in BRCA

Mutations in ACE2 gene were rare (0.30%, Figure S3A), so the mutations seemed to not be a dominating factor for ACE2 expression. The CNV pattern of ACE2 was shown in Figure S3B. Remarkably, copy number amplification of the ACE2 upregulated the expression of ACE2 mRNA (Figure S3B). Besides, methylation level was positively correlated with ACE2 expression (Figure S3C). These findings suggest that epigenetic modifications of the ACE2 gene might be essential for the regulation of expression.

Considering that ACE2 expression was not related to survival outcome in the TCGA database, we next assess its prognostic role using PrognoScan tool. However, the prognostic role of ACE2 in BRCA was inconsistent (Table S3). We speculated that the prognostic value of ACE2 may be associated with subtypes and therapeutic regimens in BRCA, which needed to be further studied.

Moreover, the functions of ACE2 in BRCA was analyzed using LinkedOmics tool. GO enrichment analysis assessed the functions of ACE2 in term of three aspects, including biological processes, cellular components and molecular functions. Plentiful of statistically significant terms were found and the top 5 terms positively correlated with ACE2 expression of each analysis were retained. As shown in Table S4, the most critical terms were associated with immune-related processes. These results reveal that ACE2 may act as a critical role in regulating anti-tumor immunity in BRCA.

### ACE2 shapes an inflamed TME in BRCA

Considering that ACE2 was positively related to a great proportion of immunomodulators in BRCA, we next explored in-depth immunological role of ACE2 in BRCA. Most MHC molecules were upregulated in the high ACE2 group, which suggested that the ability of antigen presentation and processing was upregulated in the high ACE2 group (Figure 2A). Besides, most chemokines and paired receptors were upregulated in the high ACE2 group (Figure 2A). These chemokines and receptors facilitate the recruitment of effector TIICs, including CD8+T cells, TH17 cells and antigen-presenting cells. Next, we calculated the infiltration level of TIICs using five independent algorithms. Similar to the previous results, ACE2 was positively related to the infiltration levels of the majority of immune cells using various algorithms (Figure 2B). The ESTIMATE method was next applied to estimate Tumor Purity, ESTIMATE Score, Immune Score and Stromal Score. Compared with the low ACE2 group, the high ACE2 group had enhanced ESTIMATE Score, Immune Score and Stromal Score but decreased Tumor Purity (Figure 2C). In addition, ACE2 was negatively correlated with the marker genes of immune cells, including CD8+T cell, dendritic cell, macrophage, NK cell and Th1 cell (Figure 2D). Besides, the activities of the cancer immunity cycle are a direct integrated manifestation of the functions of the chemokines and other immunomodulators. In the high ACE2 group, activities of the most steps in the cycle were revealed to be upregulated (Figure 2E). The expression of immune checkpoints such as PD-1/PD-L1 was uncovered to be high in the inflamed TME [32]. In our research, ACE2 was suggested to be positively related to most immune checkpoints, including VTCN1, PD-L1, PD-1, CTLA4 and so on (Figure 2F). Totally, ACE2 is tightly correlated with the development of an inflamed TME, which may act as a critical role in identifying the immunogenicity of BRCA.

**Figure 2.**
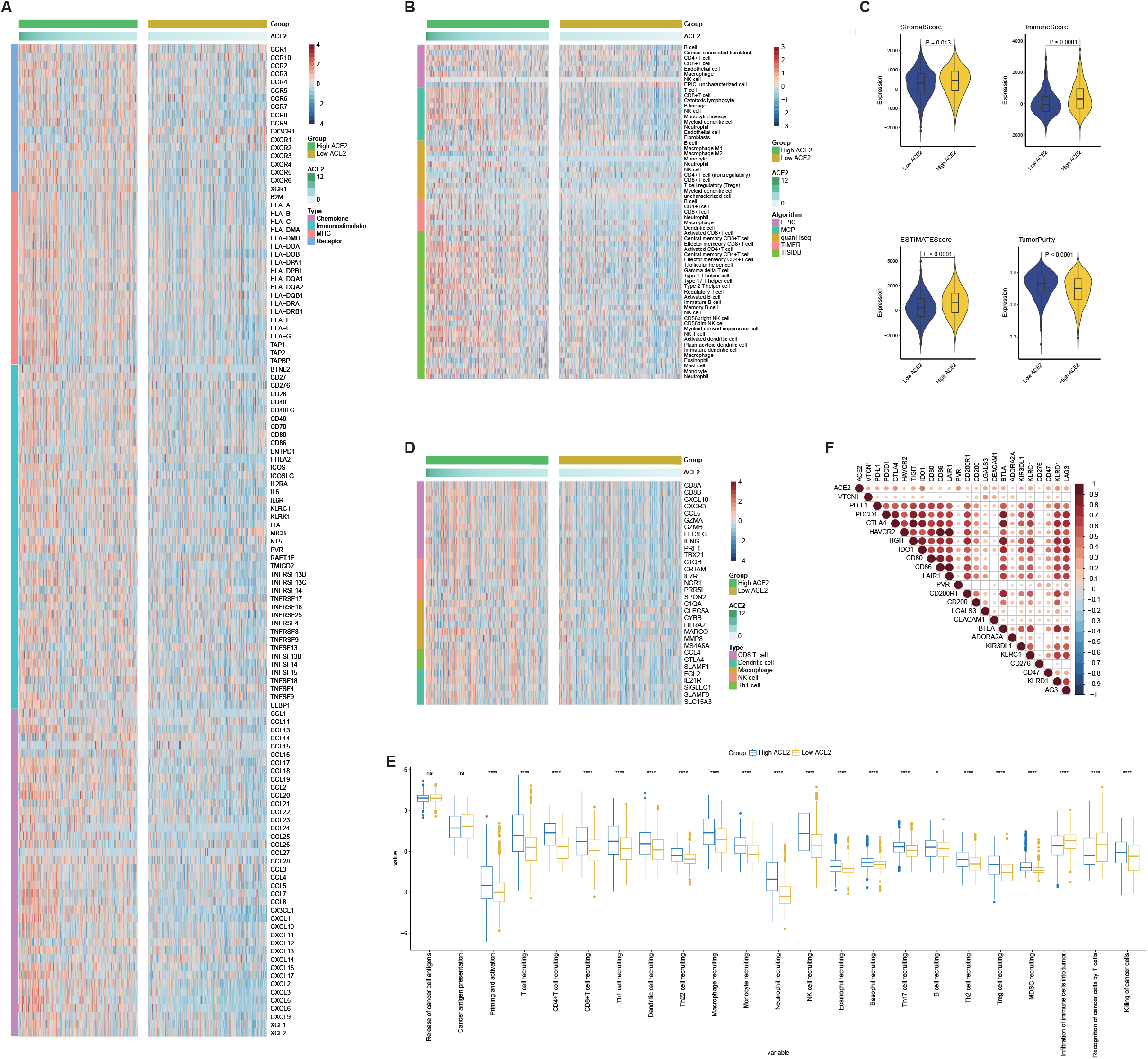
ACE2 shapes an inflamed TME in BRCA (A) Expression levels of 122 immunomodulators between the high and low ACE2 groups in BRCA. (B) Differences in the levels of TIICs calculated using five algorithms between the high and low ACE2 groups. (C) Differences in Tumor Purity, ESTIMATE Score, Immune Score, and Stromal Score estimating by ESTIMATE method between the high and low ACE2 groups. (D) Differences in the gene markers of the common TIICs between the high and low ACE2 groups. (E) Correlation between ACE2 and common inhibitory immune checkpoints. The color reveals the Pearson correlation coefficient. (F) Differences in the various steps of the cancer immunity cycle between the high and low ACE2 groups. Ns: no significant difference; *P < 0.05; **P < 0.01; ***P < 0.001; ****P < 0.0001.

### ACE2 predicts immune phenotype in BRCA

Theoretically, BRCA patients with high ACE2 expression should have a higher response to immunotherapy because ACE2 identifies an inflamed TME. Immune-related target expression levels commonly reflect the response to immunotherapy. As expected, the expression levels of most immunotargets such as CD19, PD-1 and PD-L1, were upregulated in the high ACE2 group (Figure 3A). T cell inflamed score is developed using IFN-γ-related mRNA profiles to predict clinical response to PD-1 blockade [27], and BRCA patients in the high ACE2 group exhibited higher T cell inflamed scores (Figure 3B). Besides, TMB level is another biomarker for the prediction of the response to immunotherapy [33]. Foreseeably, in the low ACE2 group, the frequency of mutant genes and TMB were both lower compared with the high ACE2 group (Figure 3C-E). Given that the TMB levels were most enriched in the range of 0-1200 (Figure S4), the comparison between the two groups was limited to the range of 1-1200 to avoid the effect of extremum. More importantly, TP53 exhibited incredibly high mutation rates in the high ACE2 group (Figure 3C-D), which was a biomarker for better response to immunotherapy [34, 35]. According to previous research, patients with high levels of microsatellite instability (MSI-H) tend to be sensitive to immunotherapy [33]. We next assess the status of mismatch repair (MMR) proteins and ACE2 expression. The proportion of MSI-H in BRCA varied largely, from 0.2% to 18.6% [36], but in most the proportion of MSI-H in BRCA less than 5%. We set the low 5% as the threshold of MMR protein deficiency. As expected, the proportion of MLH1 and PMS2 deficiency in the high ACE2 group was higher than that in the low ACE2 group, which indicated that BRCA patients with high ACE2 expression may have a higher MSI-H proportion (Figure 3F). Using IPS as a surrogate of the response to immunotherapy, we discovered that patients with high ACE2 expression had notably higher IPS (Figure 3G). In summary, immunotherapy may be carried out in BRCA patients with high ACE2 expression as they tend to be sensitive to immunotherapy.

**Figure 3.**
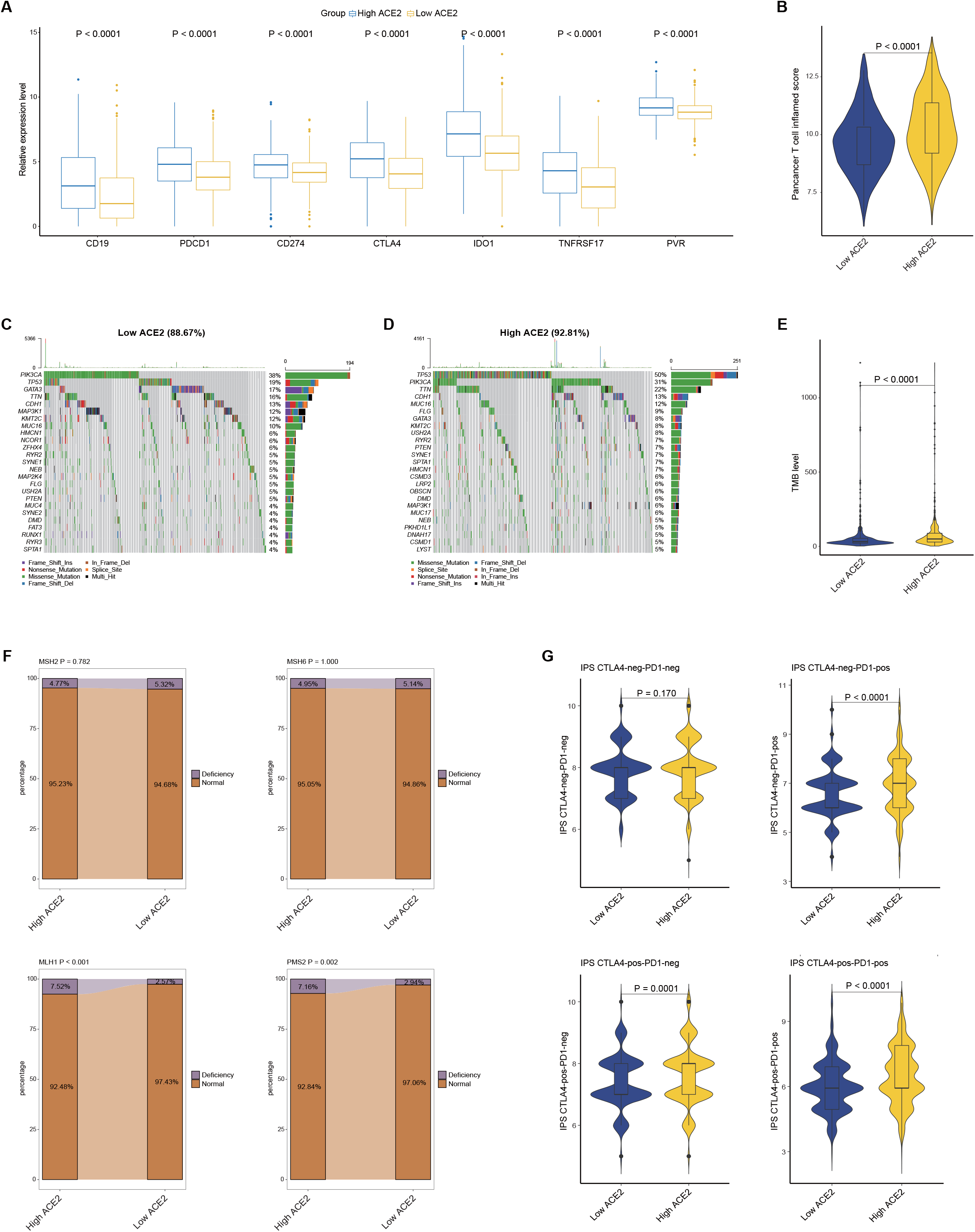
Correlation between ACE2 and the immune phenotype in BRCA (A) Differences in expression levels of immune-related targets the high and low ACE2 groups in BRCA. (B) Differences in T cell inflamed scores between the high and low ACE2 groups. The T cell inflamed score is positively related to the response to cancer immunotherapy. (C, D) Mutational landscape in the high and low ACE2 groups. (E) Differences in TMB levels between the high and low ACE2 groups. (F) Differences in deficiency rates of MMR proteins between the high and low ACE2 groups. (G) Differences in IPS scores between the high and low ACE2 groups.

### ACE2 predicts molecular subtypes and therapeutic options in BRCA

We next evaluated the ACE2 expression and clinic-pathologic features of BRCA. As Figure 4A exhibited, ACE2 expression was significantly associated with age, histological type, molecular type, ER status and PR status, but not related to other features. Specifically, ACE2 was upregulated in ER-negative, PR-negative and the triple-negative breast cancer (TNBC) tissues (Figure S5A), and ROC analysis indicated a notable diagnostic value in identifying these molecular subtypes (Figure S5B). TNBC has been identified as a subtype with high aggressiveness and PD-L1 expression. However, in the TCGA cohort, the prognosis of TNBC patients showed no notable difference compared with non-TNBC patients (Figure S6), which may be due to various therapies. This may account for why ACE2 was upregulated in TNBC subtype but not related to prognosis.

**Figure 4.**
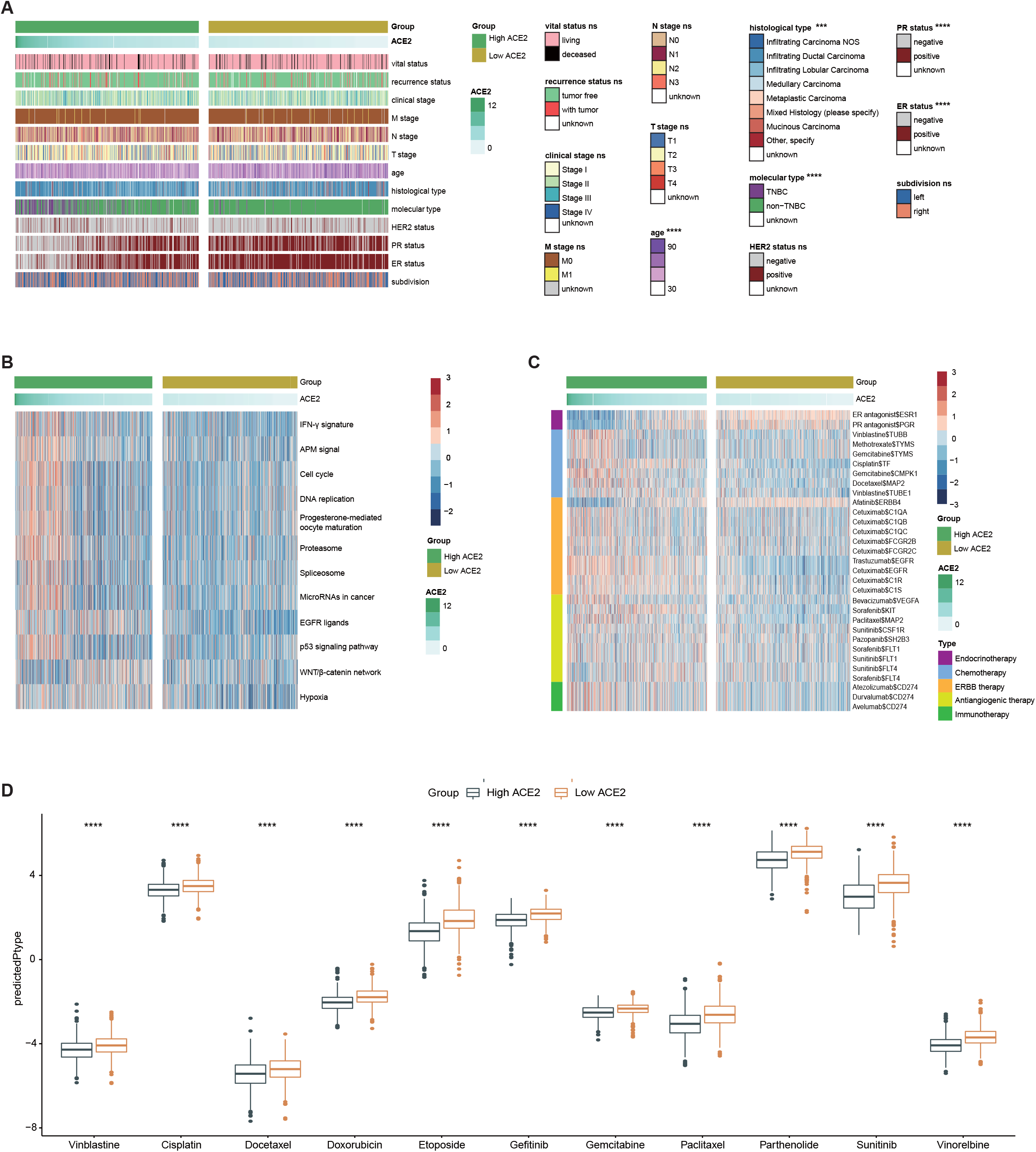
ACE2 predicts the molecular subtype and the therapeutic options in BRCA (A) Correlations between ACE2 and clinic-pathological features in BRCA. (B) Correlations between ACE2 and the enrichment scores of several therapeutic signatures. (C) Correlation between ACE2 and the drug-target genes extracted from the Drugbank database. (D) Differences in IC50 of common anti-cancer drugs between the high and low ACE2 groups. ****P < 0.0001.

In addition, the enrichment scores, such as IFN-γ signature, APM signal, cell cycle, DNA replication and et al., were higher in the high ACE2 group (Figure 4B). Thus, targeted therapy suppressing these oncogenic pathways could be applied for the treatment of BRCA with high ACE2 expression. Moreover, findings from the Drugbank database revealed a remarkably higher response to chemotherapy, anti-ERBB therapy (excluding Afatinib), antiangiogenic therapy and immunotherapy in the high ACE2 group (Figure 4C). This shows that chemotherapy, anti-ERBB therapy, antiangiogenic therapy and immunotherapy can be applied, either alone or in combination, for the therapy of BRCA with high expression. However, BRCA with lower ACE2 expression was possibly sensitive to endocrine therapy and Afatinib. Moreover, IC50 of anti-cancer drugs in patients from the TCGA database according to the pRRophetic algorithm was estimated. The results showed patients with high ACE2 expression were more sensitive to common anti-cancer drugs (Figure 4D). To sum up, ACE2 is an indicator for the subtype of TNBC and resistance to endocrine therapy in BRCA, but patients with high ACE2 expression tend to be sensitive to more therapeutic opportunities, including chemotherapy, anti-ERBB therapy, antiangiogenic therapy and immunotherapy.

### ACE2 predicts immune phenotypes and clinical subtypes in independent cohorts

The above findings were next validated in the METABRIC database. Similar to the evidence from the TCGA database, the infiltration levels of the majority of immune cells using various algorithms were increased in the high ACE2 group (Figure 5A). Besides, the enrichment scores of chemokine immunomodulator, MHC and receptor were also higher in the high ACE2 group (Figure 5B). ACE2 was positively related to the marker genes of immune cells (Figure 5C). As expected, immunotargets and T cell inflamed scores were increased in the high ACE2 group as well (Figure 5D-E). We also analyzed the association between TMB level and ACE2 expression. Although the TMB levels were not remarkably different in the two groups (Figure S7), the notably various mutant feature of TP53 was observed in the METABRIC database (Figure 5F-G). The deficient frequency of MLH1 was higher, which implied the MSI-H may be common in the high ACE2 group (Figure S8). Besides, ROC analysis validated the notable diagnostic value in identifying ER, PR status and TNBC subtype (Figure 5H). However, similar to the result in the TCGA database, the prognosis of TNBC patients showed no remarkable difference compared with non-TNBC patients (Figure S9). Furthermore, we also performed IC50 prediction of anti-cancer drugs in patients from the METABIRC database using the pRRophetic algorithm. As expected, patients with high ACE2 expression were more sensitive to these anti-cancer drugs which were mentioned in the previous analysis (Figure 5I).

**Figure 5.**
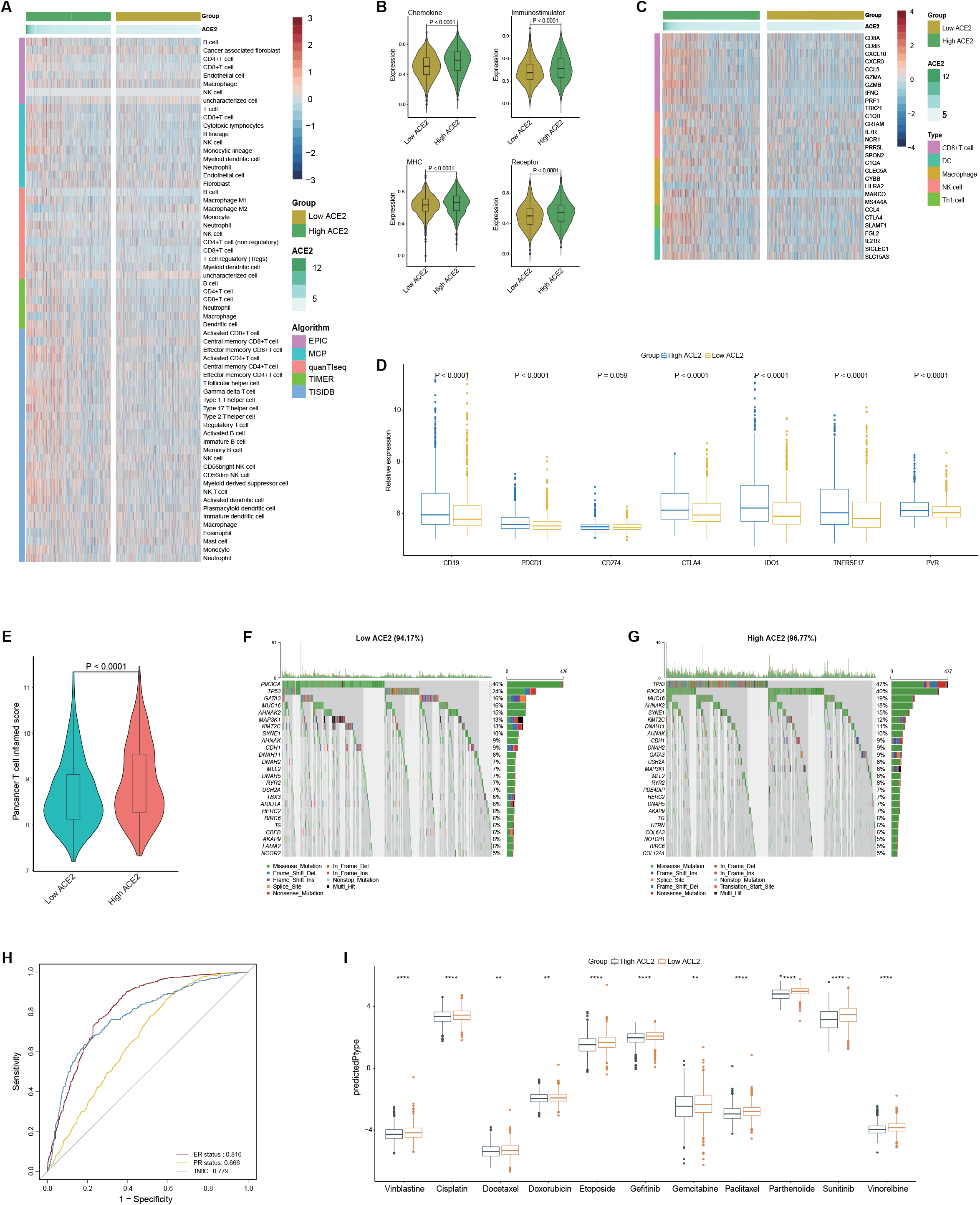
Roles of ACE2 in predicting immune and clinical phenotypes in the METABIRC cohort (A) Differences in the levels of TIICs calculated using five algorithms between the high and low ACE2 groups in BRCA. (B) Differences in enriched scores of chemokines, receptors, MHC and immunostimulators between the high and low ACE2 groups. (C) Differences in the gene markers of the common TIICs between the high and low ACE2 groups. (D) Differences in expression levels of immune-related targets between the high and low ACE2 groups. (E) Differences in T cell inflamed scores between the high and low ACE2 groups. (F, G) Mutational landscape in the high and low ACE2 groups. (H) Diagnostic values of ACE2 expression in identifying ER, PR and triple-negative subtypes. (I) Differences in IC50 of common anti-cancer drugs between the high and low ACE2 groups. **P < 0.01; ****P < 0.0001.

To further validate above results, we also obtained a TMA cohort for verification, which included 125 BRCA samples and 15 adjacent samples. As Figure 6A shown, ACE2 was notably decreased in BRCA tissues in comparison to normal tissues (Figure 6A-B). In accord with the above results, ACE2 was overexpressed in ER-negative, PR-negative and the TNBC tissues (Figure 6C). In addition, the current BRCA cohort was divided into the low and high expression groups according to the median level of ACE2 expression (IRS ≤ 3 vs. IRS > 3). As Table 1 exhibited, ACE2 expression was associated with age, ER status, PR status and molecular type (Table 1). Moreover, the infiltrating level of CD8+T cell and PD-L1 expression were higher in the high ACE2 group (Figure 6D-F). In conclusion, ACE2 expression is related to clinical features and immune phenotypes in BRCA. However, due to unavailable therapeutic information, we are unable to assess the association between ACE2 expression and the response to various therapies.

**Figure 6.**
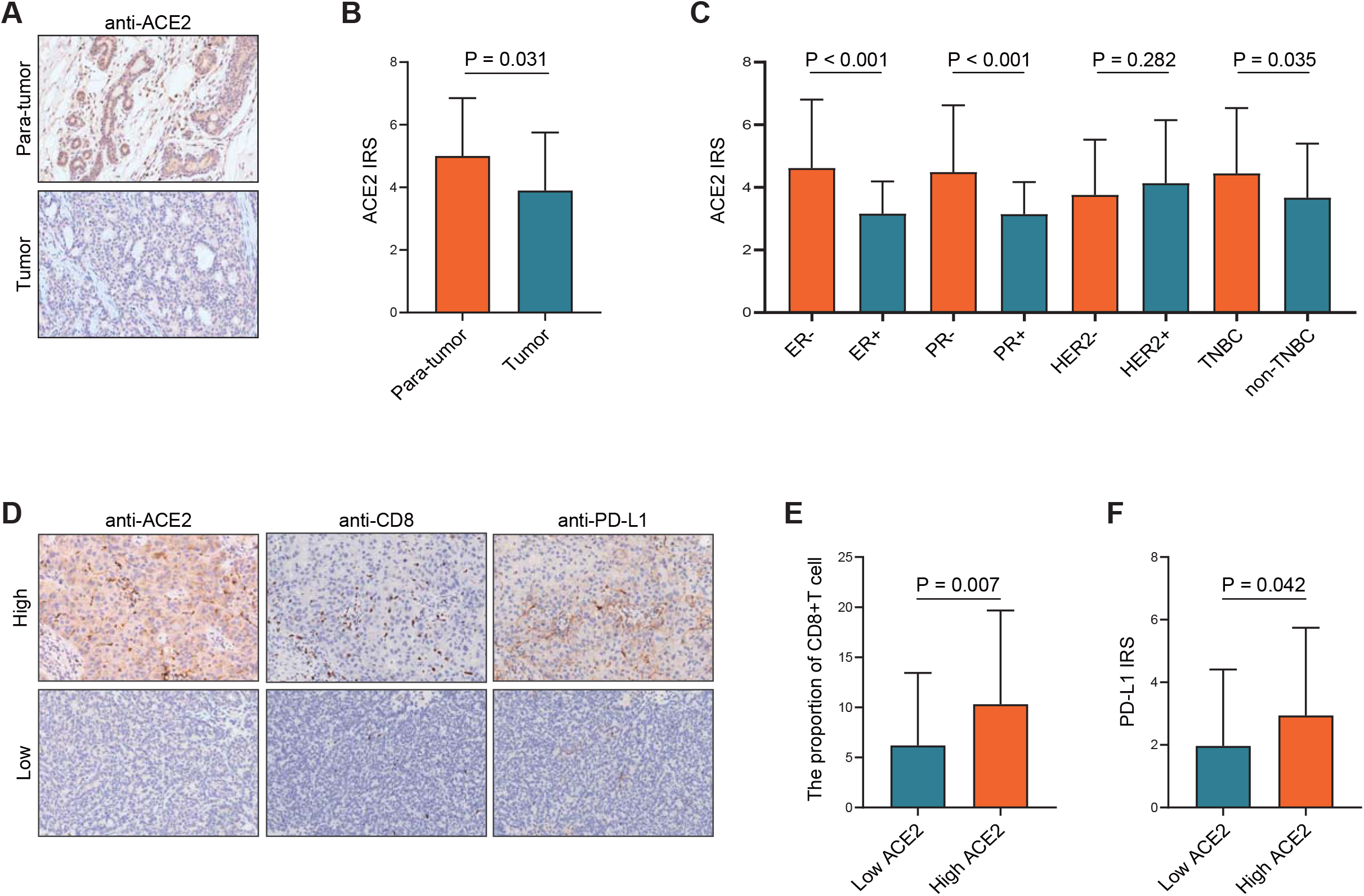
Roles of ACE2 in predicting clinical and immune phenotypes in the recruited TMA cohort (A) Representative images revealing ACE2 expression in tumor and para-tumor tissues using anti-ACE2 staining. Magnification, 200X; (B) Expression levels of ACE2 in tumor and para-tumor tissues. (C) Expression levels of ACE2 in various molecular subtypes. (D) Representative images revealing CD8+T cell infiltration and PD-L1 expression in the high and low ACE2 groups. Magnification, 200X; (E) Differences in CD8+T cell infiltration between the high and low ACE2 groups. (F) Differences in PD-L1 expression between the high and low ACE2 groups.

**Table 1.** Associations between ACE2 expression and clinico-pathological features in BRCA.

| Features | Cases | ACE2 expression | | $\chi^2$ value | P value |
| --- | --- | --- | --- | --- | --- |
|  |  | low | high |  |  |
| Age |  |  |  |  |  |
| ≤60 | 94 | 48 | 46 | 4.561 | 0.033 |
| >60 | 31 | 9 | 22 |  |  |
| T stage |  |  |  |  |  |
| ≤2cm | 47 | 18 | 29 | 1.979 | 0.159 |
| >2cm | 76 | 39 | 37 |  |  |
| unknown | 2<sup>a</sup> |  |  |  |  |
| N stage |  |  |  |  |  |
| N0 | 63 | 27 | 36 | 0.275 | 0.600 |
| N1-N3 | 61 | 29 | 32 |  |  |
| unknown | 1<sup>b</sup> |  |  |  |  |
| M stage |  |  |  |  |  |
| N0 | 124 | 56 | 68 | 1.203 | 0.456<sup>c</sup> |
| M1 | 1 | 1 | 0 |  |  |
| Clinical stage |  |  |  |  |  |
| 0-2 | 89 | 41 | 48 | 0.027 | 0.869 |
| 3-4 | 36 | 16 | 20 |  |  |
| Grade |  |  |  |  |  |
| I-II | 53 | 28 | 25 | 1.755 | 0.185 |
| II-III | 71 | 29 | 42 |  |  |
| unknown | 1 |  |  |  |  |
| ER status |  |  |  |  |  |
| negative | 63 | 18 | 45 | 14.847 | <0.001 |
| positive | 62 | 39 | 23 |  |  |
| PR status |  |  |  |  |  |
| negative | 70 | 21 | 49 | 15.607 | <0.001 |
| positive | 55 | 36 | 19 |  |  |
| HER2 status |  |  |  |  |  |
| negative | 78 | 35 | 43 | 0.102 | 0.750 |
| positive | 46 | 22 | 24 |  |  |
| unknown | 1 |  |  |  |  |
| Molecular type |  |  |  |  |  |
| non-TNBC | 88 | 47 | 41 | 6.758 | 0.009 |
| TNBC | 36 | 10 | 26 |  |  |
| unknown | 1 |  |  |  |  |

Note: a: T1-T3, not T4; b: The patient had distant metastasis; c: P value was calculated by Fisher test.

## Discussion

COVID-19 is becoming a global concern and the major public threat in the last two years. Cancer patients are more impressionable to COVID-19 infection due to the underlying disease, which may be due to systemic reduced immunity or anticancer therapy [37]. The situation could be more terrible in lung cancer patients as they already have chronic pulmonary inflammation and the lung TME supports for SARS-CoV-2 and accelerates infection [38]. Besides, cancer patients have a worse prognosis after SARS-CoV-2 infection. For example, lung cancer patients were reported to suffer more from severe events, including increased death rate and ICU admission rate [39]. Moreover, emerging studies suggest that SARS-CoV-2 infection could affect cancer progression. Dormant cancer cells tend to survive after successful therapy of primary tumors and localize in particular microanatomical sites of metastasis-prone organs [40]. Acute lung inflammation and neutrophil extracellular traps have been exhibited to activate the exit from dormancy of breast dormant cancer cells respectively, leading to distant metastasis [41].

However, an isolated case report has revealed that SARS-CoV-2 infection induced complete spontaneous remission in a patient with lymphoma [42]. This may be because of the SARS‐CoV‐2 infection activating an anti‐tumor immune response, as has been previously reported, concurrent infections induced complete remission of diffuse large B-cell lymphoma independent of any interventions [43]. Recently, oncolytic virotherapy has developed as an encouraging anti-cancer therapy through virus self-replication. On the other side, oncolytic virus enhances anti-tumor immunity responses by increasing the immune infiltration and turn the tumor to be sensitive to immunotherapy and chemotherapy [44]. Thus, we speculated SARS‐CoV‐2 infection induced spontaneous remission of tumor has parallels with oncolytic virotherapy to some extent. Whatever, the complicated interplay between the immune system and cancer after SARS‐CoV‐2 infection needs to be further highlighted.

As crosstalk between COVID-19 and the tumor, ACE2 act as a significant role in both SARS‐CoV‐2 infection and cancer. ACE2 has been found to be as a tumor suppressor in various cancers. It was reported that ACE2 exerts anti-tumor roles by suppressing tumor angiogenesis [8]. Dai et al. [11] reported that upregulated ACE2 expression was correlated with a favorable clinical outcome in hepatocellular carcinoma. Besides, the associations between ACE2 with anti-tumor immunity and immunotherapy were also explored. Yang et al. [12] reported that the overexpression of ACE2 was significantly associated with enhanced immune infiltration in endometrial cancer and renal papillary cell cancer. Bao et al. [31] uncovered tight correlations between ACE2 expression and immune gene signatures in multiple cancers. However, the crucial values of ACE2 in the identification of tumor immune status have not been evaluated.

In this research, we first conducted a pan-cancer analysis of the immunological features of ACE2. We discovered that ACE2 exhibited the tightest correlation with immunological factors in BRCA, and in-depth analysis of ACE2 in BRCA was subsequently conducted. We found that ACE2 was positively related to the expression of important immunomodulators, such as CCL5, CXCL10 and CXCR4. Besides, ACE2 was also positively related to increased TIICs and cancer immunity cycles. Namely, the recruitment of effector TIICs was enhanced, thereby facilitating the formation of an inflamed TME. Besides, we discovered that high ACE2 expression was correlated with the enhanced response to immunotherapy by check the difference of immune checkpoints expression, T cell inflamed score, TMB, MMR protein deficiency status and IPS scores in the ACE2 high and low groups. Another important finding was that high ACE2 levels indicated the negativity of hormone receptors, including ER, PR and HER2 receptors. More importantly, ACE2 expression was increased in the TNBC subtype of BRCA. TNBC is summarized by deadly aggressiveness and lacked treatment [45]. However, PD-L1 was often overexpressed in TNBC [46], and its response to immune checkpoint inhibition was encouraging.

## Conclusions

In the current study, we revealed that ACE2 shaped an inflamed TME according to the evidence that ACE2 positively related to the immunological patterns of TME in BRCA. Besides, we uncovered that BRCA had the potential to estimate the response of immunotherapy, the molecular subtypes and the response to several therapeutic strategies. Overall, ACE2 may be used as a promising biomarker for the identification of immunological features in BRCA.

## Acknowledgments

This work was supported by the Natural Science Foundation of China (82073194), the Major project of Wuxi Science and Technology Bureau (N20201006) and the Wuxi Double-Hundred Talent Fund Project (BJ2020076).

## Availability of data and materials

All data supported the results in this study are showed in this published article and its supplementary files. Besides, original data for bioinformatics analysis could be downloaded from corresponding platforms.

## Authors’ contributions

Y Zhang and Y Zhu conceived the study and participated in the study design, performance, coordination and project supervision. JM, YC, RX, XY, LC, TM, TG and FG collected the public data and performed the bioinformatics analysis. JM performed the IHC staining. Y Zhang and Y Zhu revised the manuscript. All authors reviewed and approved the final manuscript.

## Ethics approval and consent to participate

Ethical approval for the study of tissue microarray slide was granted by the Clinical Research Ethics Committee, Outdo Biotech (Shanghai, China).

## Supplemental Figure legends

**Figure S1.**
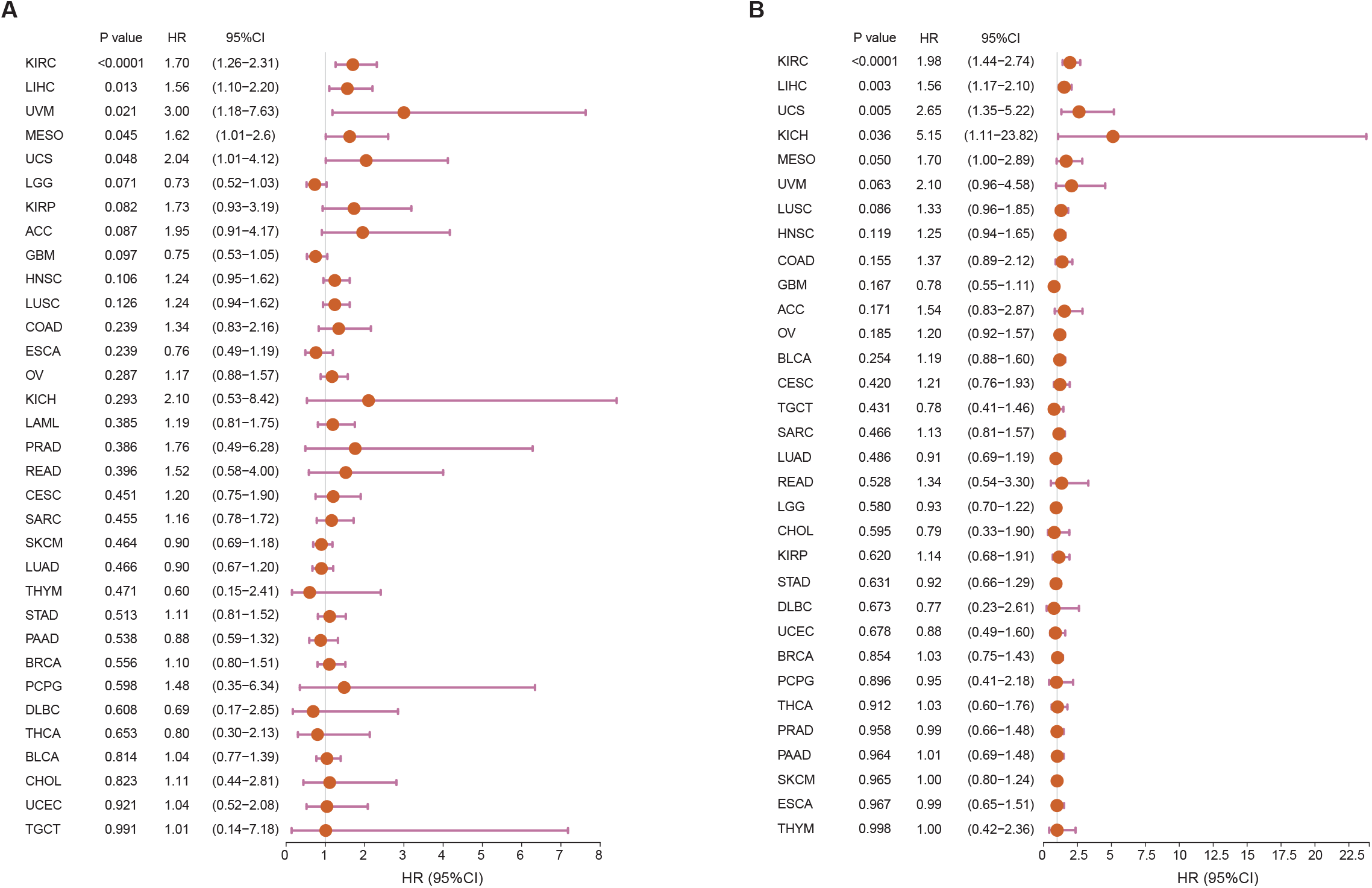
Forest plots of prognostic value of ACE2 in predicting (A) OS and (B) PFS in pan-cancer analysis.

**Figure S2.**
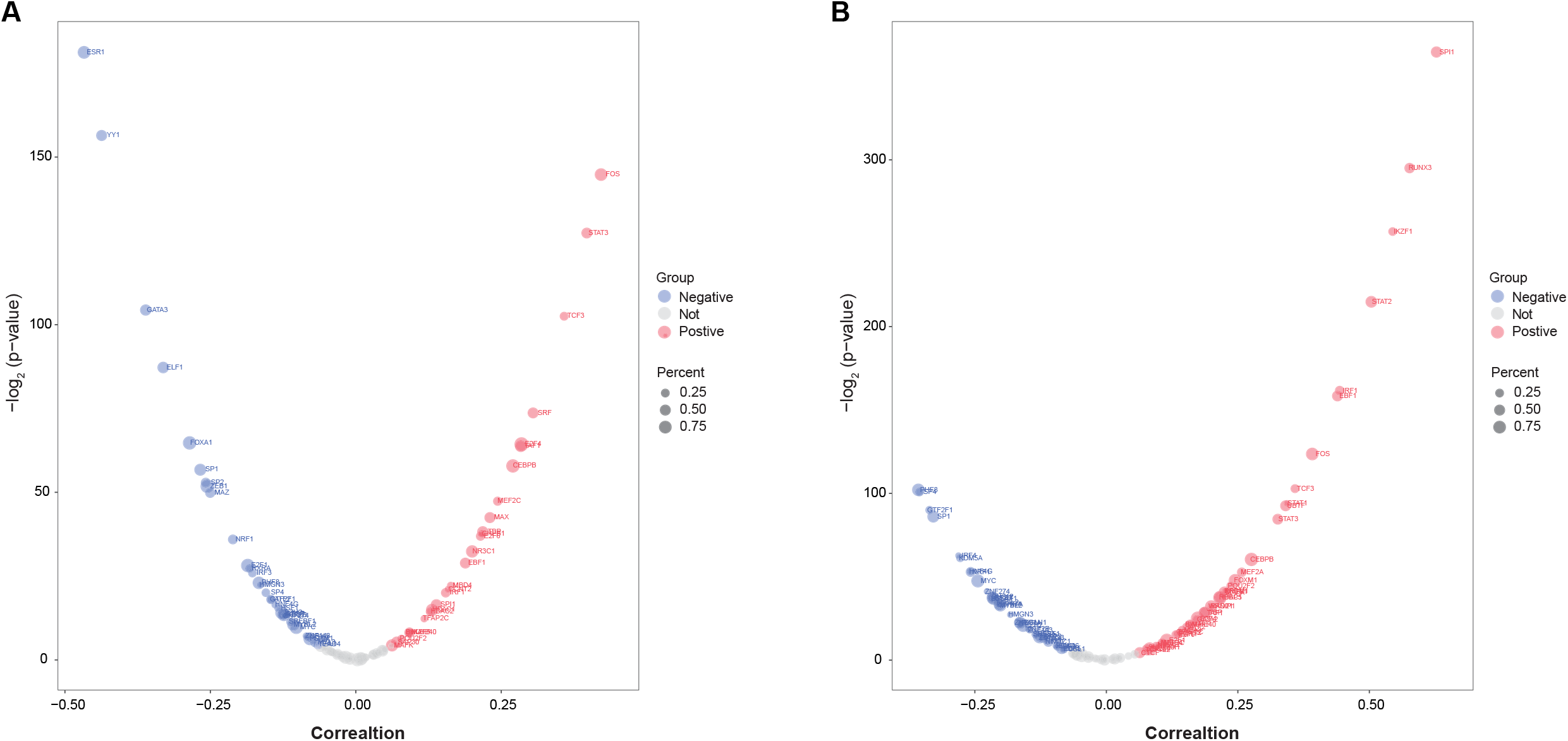
TFs that correlated with (A) ACE2 and (B) PD-L1 calculated by RABIT algorithm in BRCA.

**Figure S3.**
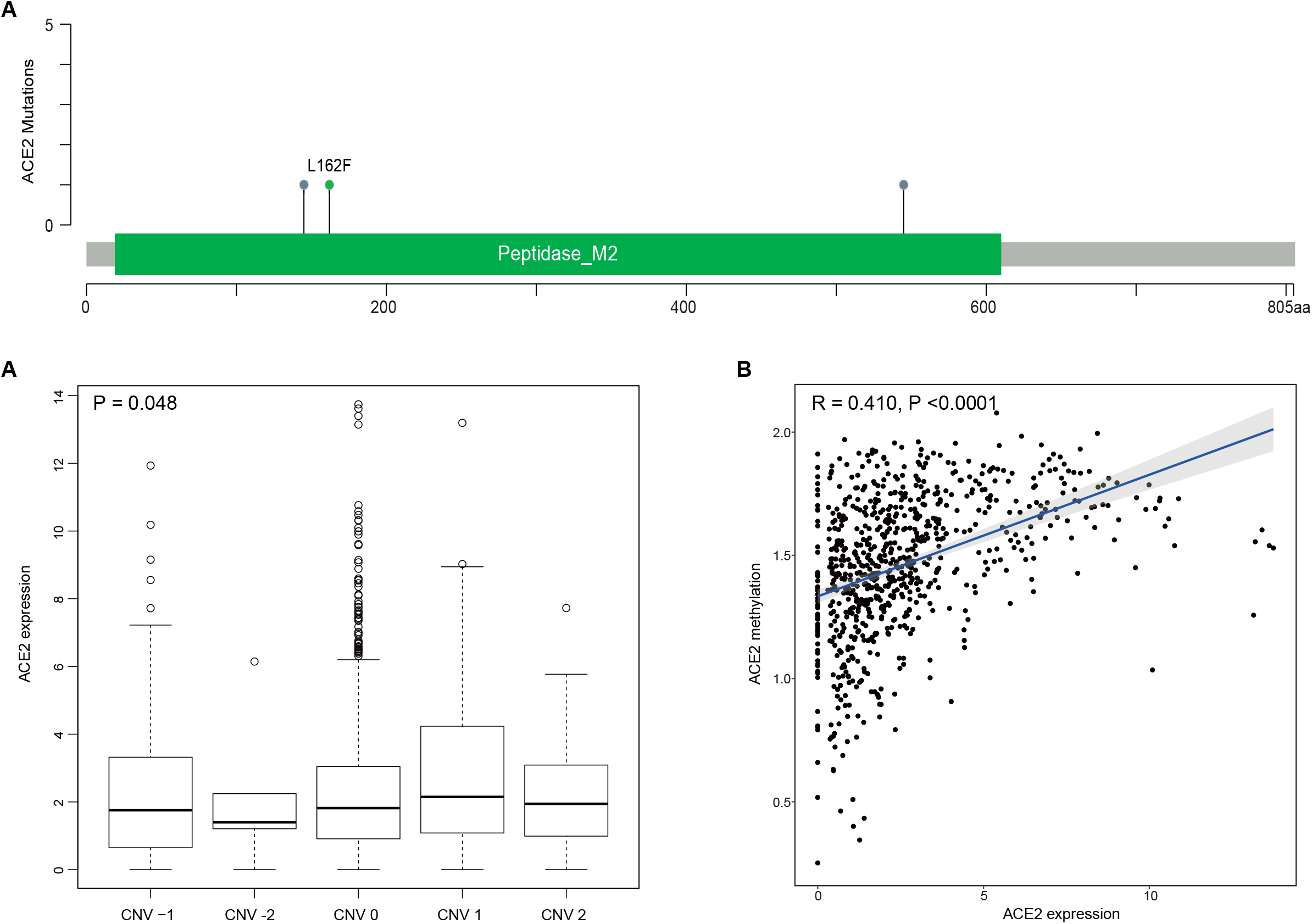
Potential regulatory factors of ACE2 in BRCA. (A) Mutations in ACE2 gene. (B) The associations between CNV pattern and ACE2 expression in BRCA. (C) The correlation between methylation level and ACE2 expression.

**Figure S4.**
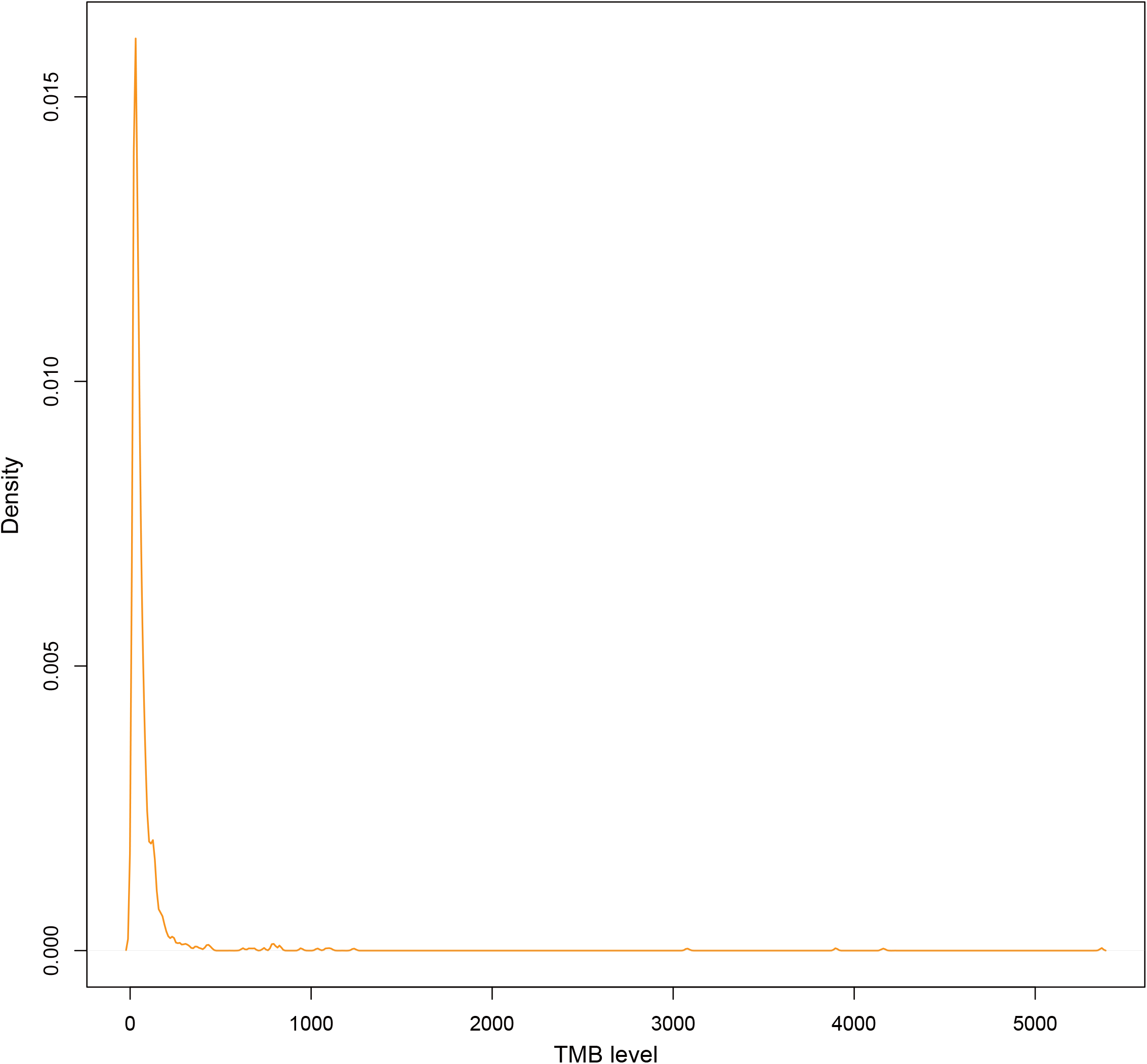
Mutational density curve in BRCA. The TMB levels were most enriched in the range of 0-1200.

**Figure S5.**
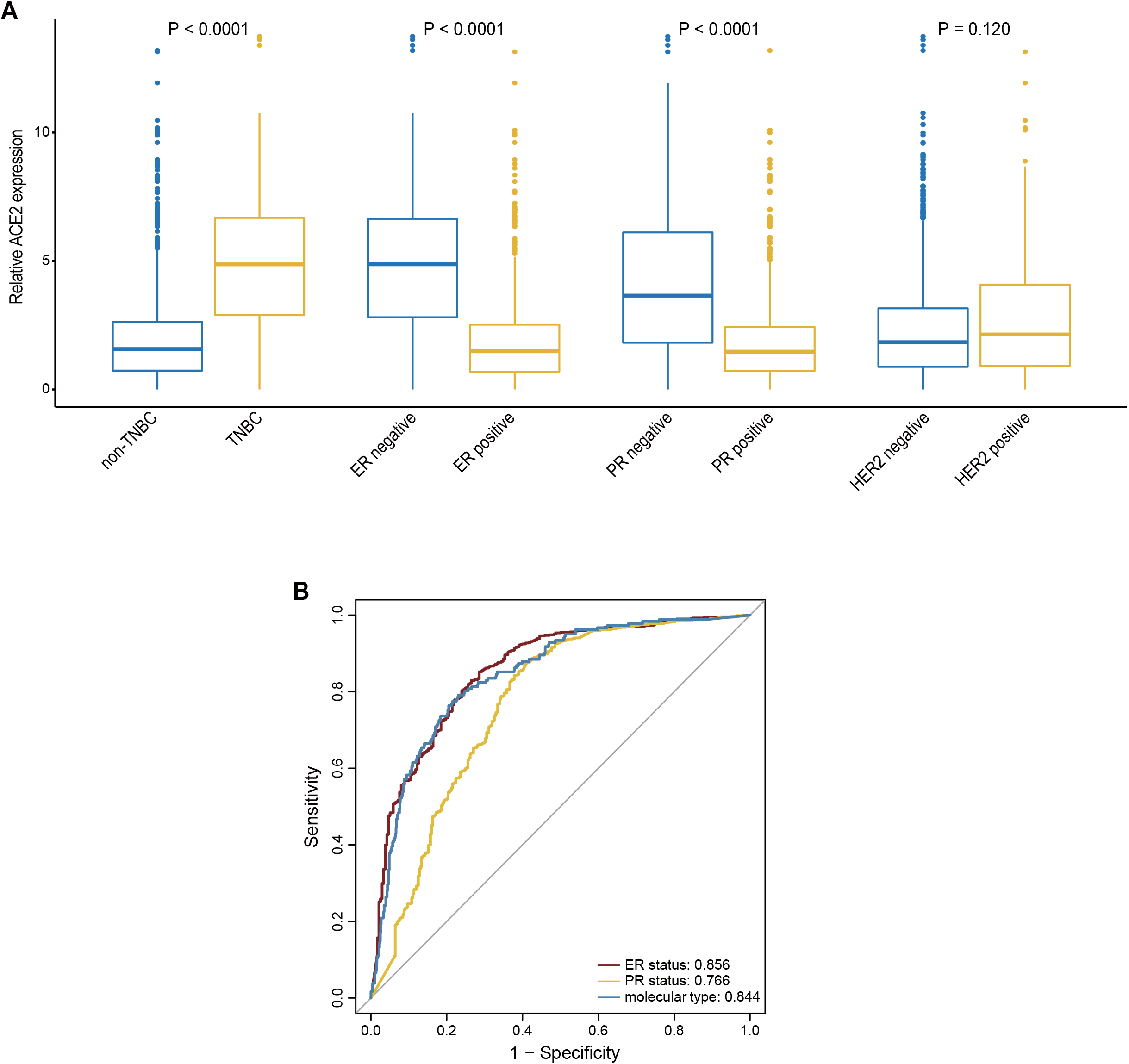
ACE2 predicts the molecular subtype in BRCA (A) Expression level of ACE2 in different molecular subtypes in BRCA. (B) Diagnostic values of ACE2 expression in identifying ER, PR and triple-negative subtypes.

**Figure S6.**
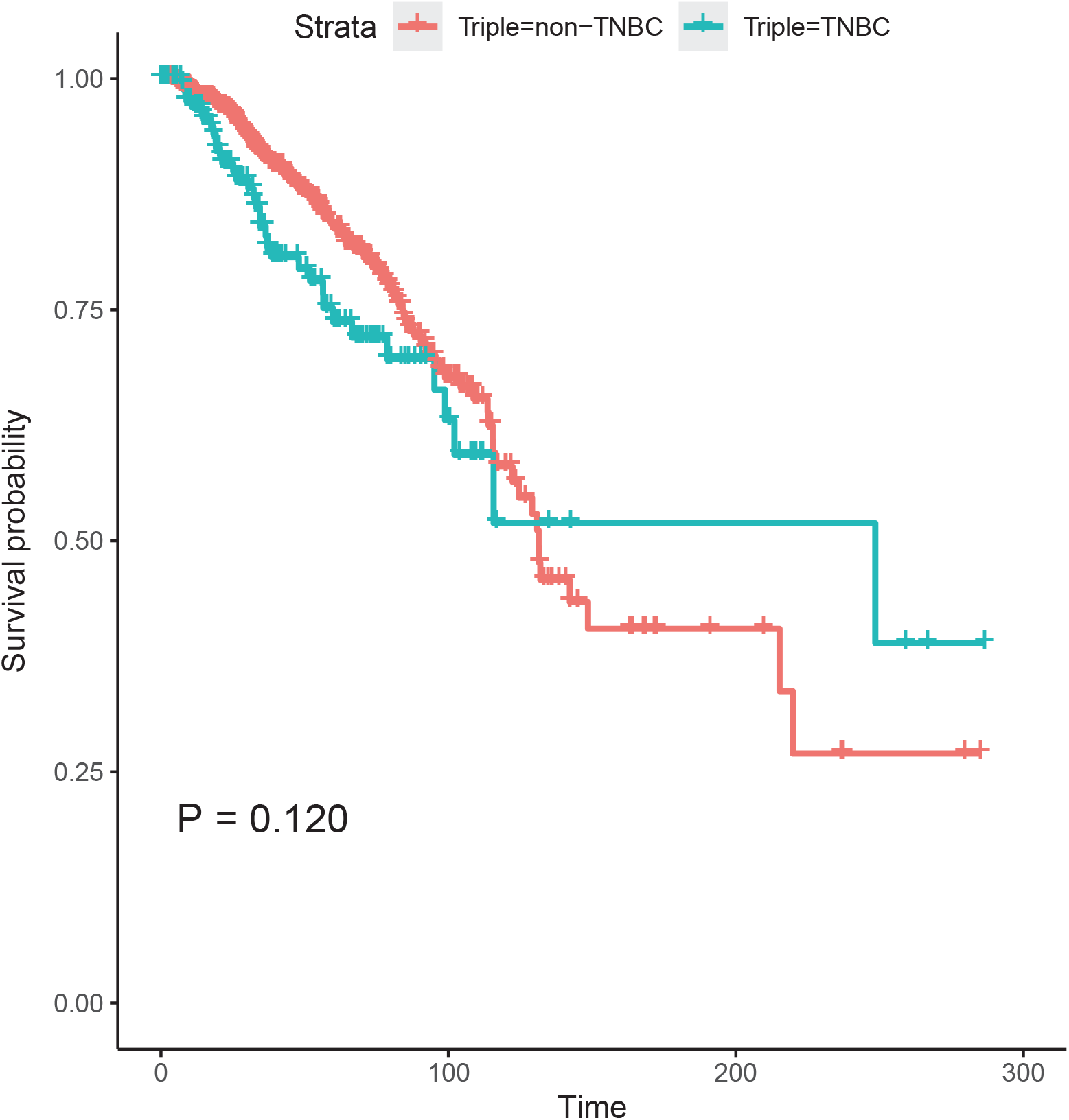
Survival analysis of patients with TNBC compared with that with non-TNBC (TCGA)

**Figure S7.**
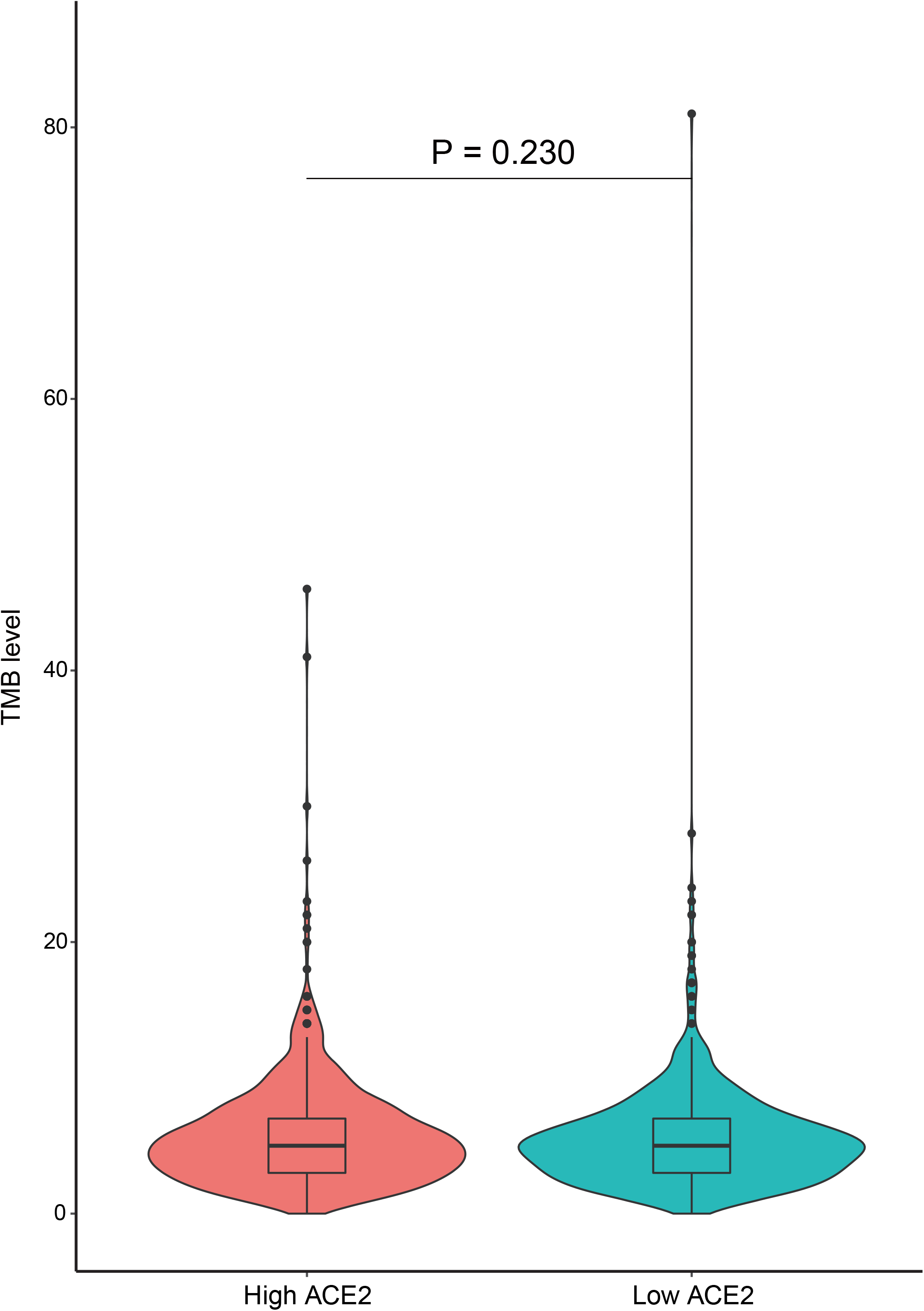
Differences in TMB levels between the high and low ACE2 groups in BRCA (METABIRC).

**Figure S8.**
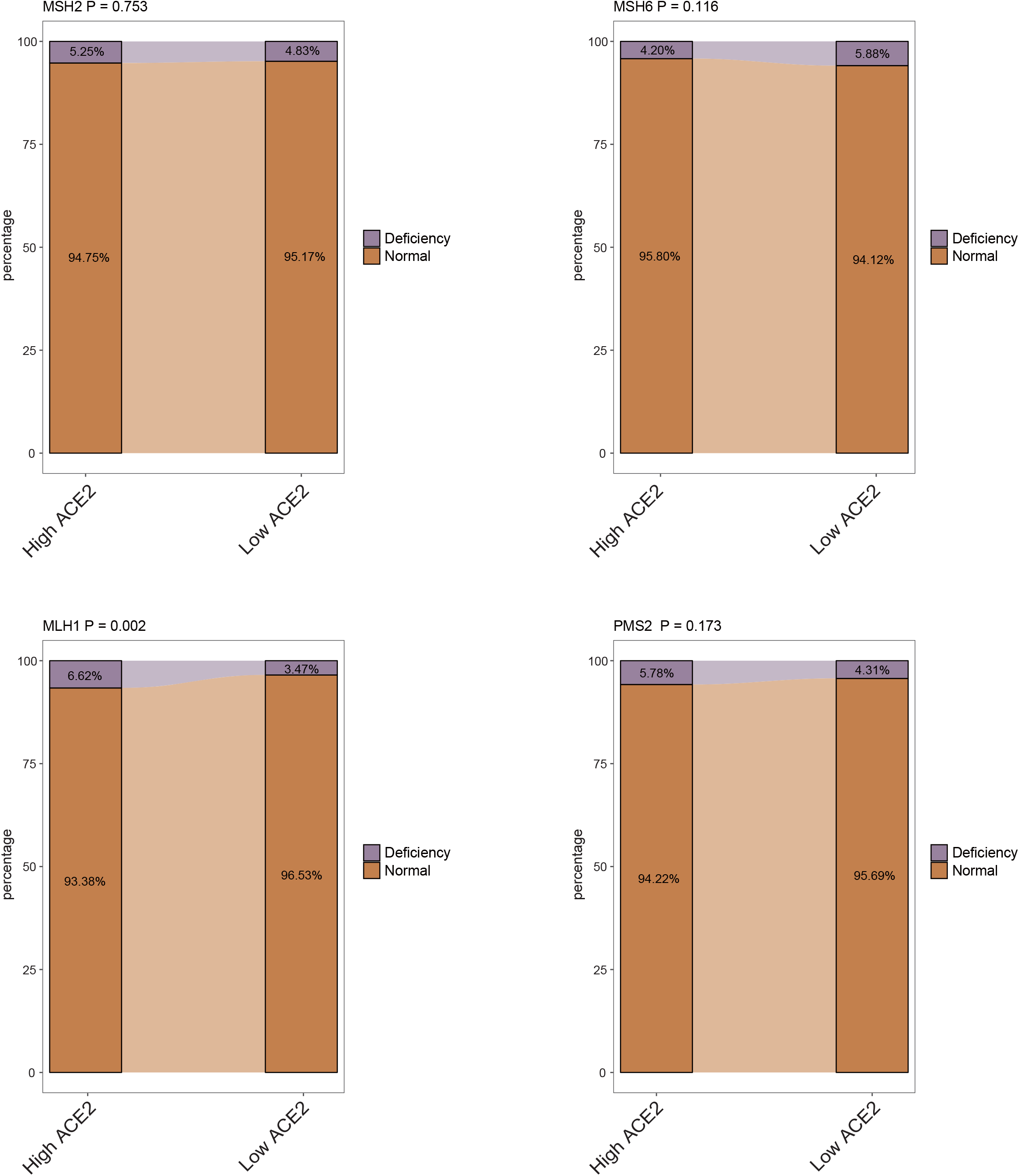
Differences in deficiency rates of MMR proteins between the two groups (METABIRC).

**Figure S9.**
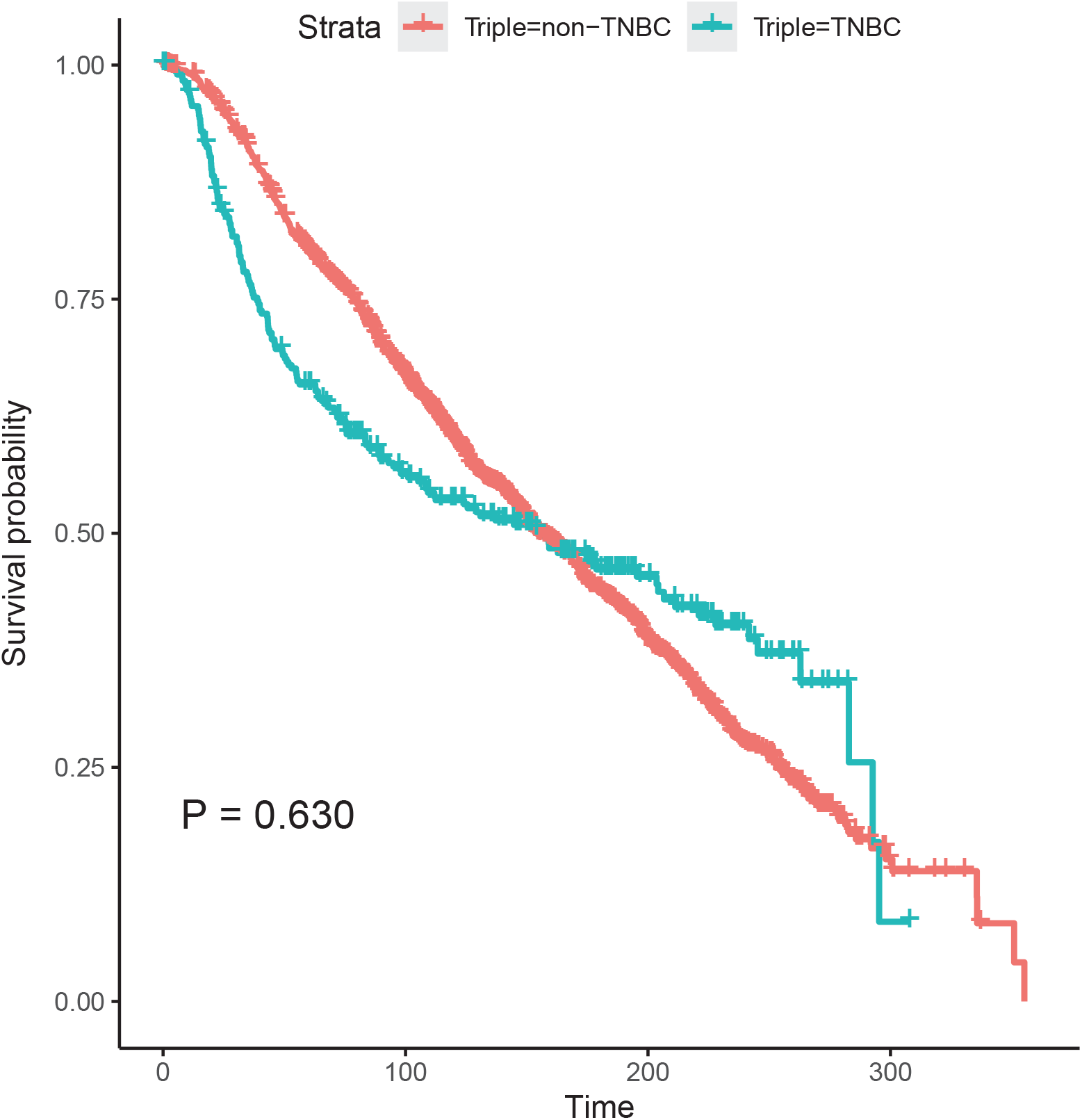
Survival analysis of patients with TNBC compared with that with non-TNBC (METABIRC).

**Table S1.** Table of abbreviations.

| Abbreviation | Full name |
| --- | --- |
| ACC | Adrenocortical carcinoma |
| BLCA | Bladder urothelial carcinoma |
| BRCA | Breast invasive carcinoma |
| CESC | Cervical squamous cell carcinoma and endocervical adenocarcinoma |
| CHOL | Cholangio carcinoma |
| COAD | Colon adenocarcinoma |
| DLBC | Lymphoid neoplasm diffuse large B-cell lymphoma |
| ESCA | Esophageal carcinoma |
| GBM | Glioblastoma multiforme |
| HNSC | Head and neck squamous cell carcinoma |
| KICH | Kidney chromophobe carcinoma |
| KIRC | Kidney renal clear cell carcinoma |
| KIRP | Kidney renal papillary cell carcinoma |
| LAML | Acute myeloid leukemia |
| LGG | Brain lower grade glioma |
| LIHC | Liver hepatocellular carcinoma |
| LUAD | Lung adenocarcinoma |
| LUSC | Lung squamous cell carcinoma |
| MESO | Mesothelioma |
| OV | Ovarian serous cystadenocarcinoma |
| PAAD | Pancreatic adenocarcinoma |
| PCPG | Pheochromocytoma and paraganglioma |
| PRAD | Prostate adenocarcinoma |
| READ | Rectum adenocarcinoma |
| SARC | Sarcoma |
| SKCM | Skin cutaneous melanoma |
| STAD | Stomach adenocarcinoma |
| TGCT | Testicular germ cell tumors |
| THCA | Thyroid carcinoma |
| THYM | Thymoma |
| UCEC | Uterine corpus endometrial carcinoma |
| UCS | Uterine carcinosarcoma |
| UVM | Uveal melanoma |

**Table S2.**
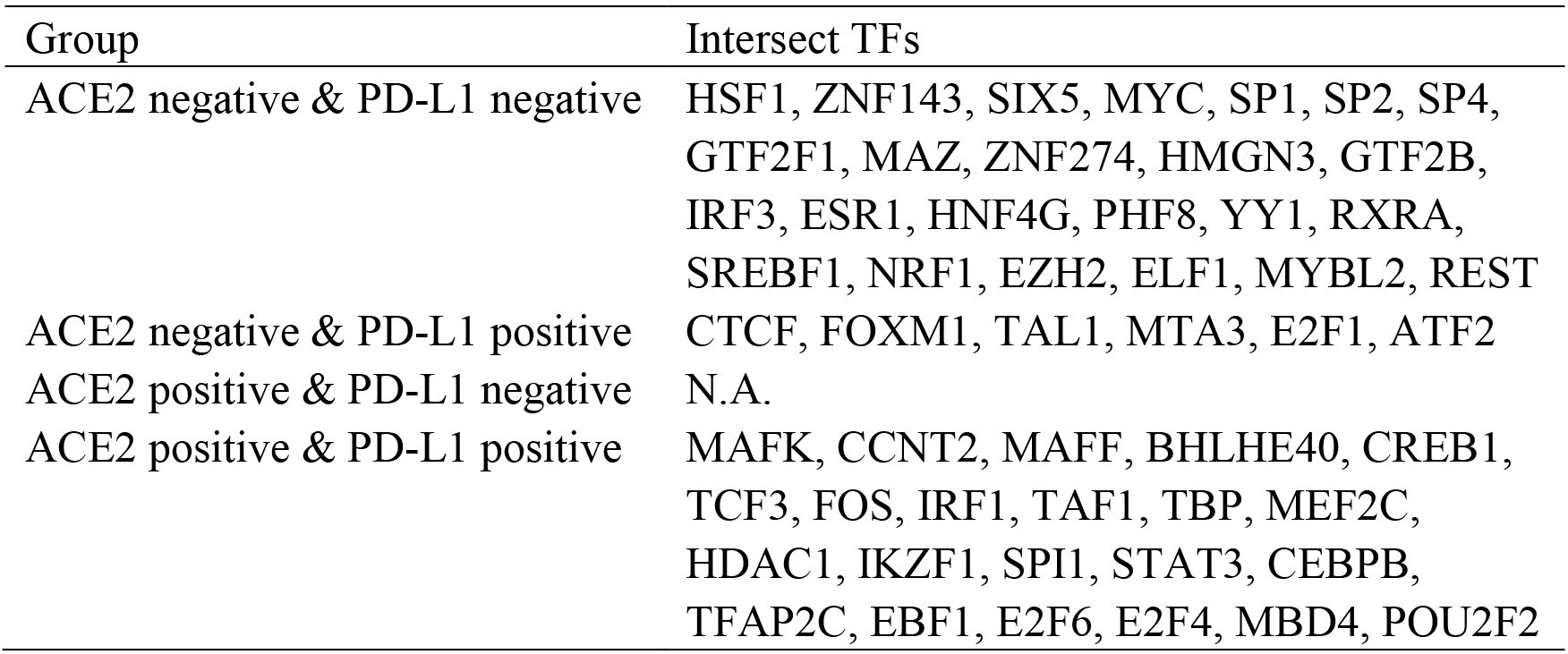
TFs that correlate with ACE2 and PD-L1.

| Group | Intersect TFs |
| --- | --- |
| ACE2 negative & PD-L1 negative | HSF1, ZNF143, SIX5, MYC, SP1, SP2, SP4, GTF2F1, MAZ, ZNF274, HMGN3, GTF2B, IRF3, ESR1, HNF4G, PHF8, YY1, RXRA, SREBF1, NRF1, EZH2, ELF1, MYBL2, REST |
| ACE2 negative & PD-L1 positive | CTCF, FOXM1, TAL1, MTA3, E2F1, ATF2 |
| ACE2 positive & PD-L1 negative | N.A. |
| ACE2 positive & PD-L1 positive | MAFK, CCNT2, MAFF, BHLHE40, CREB1, TCF3, FOS, IRF1, TAF1, TBP, MEF2C, HDAC1, IKZF1, SPI1, STAT3, CEBPB, TFAP2C, EBF1, E2F6, E2F4, MBD4, POU2F2 |

**Table S3.** Prognostic values of ACE2 in BRCA across public microarray datasets.

| Dataset | Probe ID | Cases | Endpoint | HR | 95% CI | P value |
| --- | --- | --- | --- | --- | --- | --- |
| GSE7378 | 219962_at | 54 | DFS | 0.26 | 0.01 - 8.86 | 0.175 |
| GSE7378 | 222257_s_at | 54 | DFS | 0.41 | 0.02 - 7.59 | 0.154 |
| GSE4922-GPL96 | 222257_s_at | 249 | DFS | 0.71 | 0.49 - 1.04 | 0.043 |
| GSE4922-GPL96 | 219962_at | 249 | DFS | 1.03 | 0.85 - 1.24 | 0.145 |
| GSE1456-GPL96 | 222257_s_at | 159 | DSS | 1.41 | 0.89 - 2.26 | 0.007 |
| GSE1456-GPL96 | 219962_at | 159 | DSS | 1.04 | 0.74 - 1.46 | 0.186 |
| E-TABM-158 | 222257_s_at | 117 | DSS | 1.22 | 0.66 - 2.26 | 0.124 |
| E-TABM-158 | 219962_at | 117 | DSS | 1.48 | 0.80 - 2.74 | 0.118 |
| GSE3494-GPL96 | 219962_at | 236 | DSS | 1.11 | 0.87 - 1.43 | 0.096 |
| GSE3494-GPL96 | 222257_s_at | 236 | DSS | 0.77 | 0.47 - 1.27 | 0.062 |
| GSE19615 | 219962_at | 115 | DMFS | 1.32 | 0.72 - 2.41 | 0.038 |
| GSE19615 | 222257_s_at | 115 | DMFS | 1.47 | 0.92 - 2.35 | 0.024 |
| GSE6532-GPL570 | 219962_at | 87 | DMFS | 3.76 | 0.91 - 15.59 | 0.065 |
| GSE6532-GPL570 | 222257_s_at | 87 | DMFS | 3.36 | 1.21 - 9.30 | 0.168 |
| GSE9195 | 219962_at | 77 | DMFS | 0.30 | 0.01 - 12.36 | 0.109 |
| GSE9195 | 222257_s_at | 77 | DMFS | 0.44 | 0.05 - 4.21 | 0.192 |
| GSE12093 | 219962_at | 136 | DMFS | 0.93 | 0.62 - 1.39 | 0.197 |
| GSE12093 | 222257_s_at | 136 | DMFS | 1.08 | 0.71 - 1.66 | 0.306 |
| GSE11121 | 219962_at | 200 | DMFS | 0.81 | 0.62 - 1.04 | 0.059 |
| GSE11121 | 222257_s_at | 200 | DMFS | 0.73 | 0.53 - 1.02 | 0.013 |
| GSE2034 | 219962_at | 286 | DMFS | 0.93 | 0.79 - 1.10 | 0.080 |
| GSE2034 | 222257_s_at | 286 | DMFS | 0.92 | 0.72 - 1.16 | 0.068 |
| E-TABM-158 | 219962_at | 117 | DMFS | 1.36 | 0.73 - 2.54 | 0.133 |
| E-TABM-158 | 222257_s_at | 117 | DMFS | 1.41 | 0.87 - 2.29 | 0.060 |
| GSE2990 | 219962_at | 125 | DMFS | 1.55 | 1.02 - 2.34 | 0.070 |
| GSE2990 | 222257_s_at | 125 | DMFS | 1.54 | 0.86 - 2.76 | 0.082 |
| GSE2990 | 219962_at | 54 | DMFS | 0.81 | 0.25 - 2.61 | 0.100 |
| GSE2990 | 222257_s_at | 54 | DMFS | 0.79 | 0.41 - 1.49 | 0.102 |
| GSE7390 | 222257_s_at | 198 | DMFS | 1.19 | 1.04 - 1.36 | 0.005 |
| GSE7390 | 219962_at | 198 | DMFS | 1.16 | 1.03 - 1.30 | 0.003 |
| GSE9893 | 10330 | 155 | OS | 0.83 | 0.61 - 1.11 | 0.027 |
| GSE1456-GPL96 | 219962_at | 159 | OS | 1.05 | 0.78 - 1.40 | 0.247 |
| GSE1456-GPL96 | 222257_s_at | 159 | OS | 1.28 | 0.84 - 1.93 | 0.029 |
| E-TABM-158 | 219962_at | 117 | OS | 1.39 | 0.78 - 2.47 | 0.232 |
| E-TABM-158 | 222257_s_at | 117 | OS | 1.19 | 0.69 - 2.07 | 0.204 |
| GSE7390 | 222257_s_at | 198 | OS | 1.23 | 1.08 - 1.41 | 0.001 |
| GSE7390 | 219962_at | 198 | OS | 1.18 | 1.05 - 1.33 | 0.002 |
| GSE12276 | 222257_s_at | 204 | RFS | 1.11 | 1.00 - 1.22 | 0.005 |
| GSE12276 | 219962_at | 204 | RFS | 1.12 | 1.02 - 1.22 | 0.003 |
| GSE6532-GPL570 | 219962_at | 87 | RFS | 3.76 | 0.91 - 15.59 | 0.065 |
| GSE6532-GPL570 | 222257_s_at | 87 | RFS | 3.36 | 1.21 - 9.30 | 0.168 |
| GSE9195 | 219962_at | 77 | RFS | 0.49 | 0.05 - 4.52 | 0.107 |
| GSE9195 | 222257_s_at | 77 | RFS | 0.60 | 0.13 - 2.70 | 0.130 |
| GSE1378 | 21336 | 60 | RFS | 1.38 | 1.06 - 1.79 | 0.001 |
| GSE1379 | 21336 | 60 | RFS | 1.36 | 1.08 - 1.73 | 0.000 |
| GSE1456-GPL96 | 219962_at | 159 | RFS | 1.04 | 0.77 - 1.39 | 0.239 |
| GSE1456-GPL96 | 222257_s_at | 159 | RFS | 1.19 | 0.77 - 1.83 | 0.075 |
| E-TABM-158 | 222257_s_at | 117 | RFS | 1.19 | 0.69 - 2.07 | 0.204 |
| E-TABM-158 | 219962_at | 117 | RFS | 1.39 | 0.78 - 2.47 | 0.232 |
| GSE2990 | 222257_s_at | 62 | RFS | 0.79 | 0.47 - 1.31 | 0.031 |
| GSE2990 | 219962_at | 125 | RFS | 1.35 | 0.88 - 2.06 | 0.113 |
| GSE2990 | 222257_s_at | 125 | RFS | 1.32 | 0.75 - 2.30 | 0.078 |
| GSE2990 | 219962_at | 62 | RFS | 0.71 | 0.27 - 1.86 | 0.145 |
| GSE7390 | 219962_at | 198 | RFS | 1.06 | 0.95 - 1.17 | 0.034 |
| GSE7390 | 222257_s_at | 198 | RFS | 1.09 | 0.96 - 1.24 | 0.033 |

DFS: Disease Free Survival; DSS: Disease Specific Survival; DMFS: Distant Metastasis Free Survival; OS: Overall Survival; RFS: Relapse Free Survival.

**Table S4.** GO and KEGG pathway enrichment analyses of ACE2 in BRCA.

| Gene Set | Description | ES | NES | P value |
| --- | --- | --- | --- | --- |
| Biological process |  |  |  |  |
| GO:0002250 | adaptive immune response | 0.592 | 1.974 | <0.001 |
| GO:0002237 | response to molecule of bacterial origin | 0.576 | 1.900 | <0.001 |
| GO:1990868 | response to chemokine | 0.655 | 1.895 | <0.001 |
| GO:0060326 | cell chemotaxis | 0.568 | 1.867 | <0.001 |
| GO:0031349 | positive regulation of defense response | 0.559 | 1.865 | <0.001 |
| Cell component |  |  |  |  |
| GO:0005790 | smooth endoplasmic reticulum | 0.717 | 1.749 | <0.001 |
| GO:0001533 | cornified envelope | 0.689 | 1.898 | <0.001 |
| GO:0097038 | perinuclear endoplasmic reticulum | 0.682 | 1.544 | 0.020 |
| GO:0001891 | phagocytic cup | 0.671 | 1.533 | 0.019 |
| GO:0042611 | MHC protein complex | 0.666 | 1.452 | 0.023 |
| Molecular function |  |  |  |  |
| GO:0016701 | oxidoreductase activity, acting on single donors with incorporation of molecular oxygen | 0.774 | 1.779 | <0.001 |
| GO:0016917 | GABA receptor activity | 0.755 | 1.670 | <0.001 |
| GO:0042287 | MHC protein binding | 0.725 | 1.707 | <0.001 |
| GO:0001530 | lipopolysaccharide binding | 0.720 | 1.689 | 0.003 |
| GO:0070003 | threonine-type peptidase activity | 0.688 | 1.563 | 0.024 |
| KEGG |  |  |  |  |
| hsa04650 | Natural killer cell mediated cytotoxicity | 0.654 | 1.987 | <0.001 |
| hsa04060 | Cytokine-cytokine receptor interaction | 0.599 | 1.969 | <0.001 |
| hsa05340 | Primary immunodeficiency | 0.741 | 1.889 | <0.001 |
| hsa00601 | Glycosphingolipid biosynthesis | 0.803 | 1.875 | <0.001 |
| hsa00380 | Tryptophan metabolism | 0.726 | 1.843 | <0.001 |

ES: Enrichment score; NES: Normalized enrichment score.

